# Caffeine-induced Plasticity of Grey Matter Volume in Healthy Brains: A placebo-controlled multimodal within-subject study

**DOI:** 10.1101/804047

**Authors:** Y.-S. Lin, J. Weibel, H.-P. Landolt, F. Santini, M. Meyer, S. Borgwardt, C. Cajochen, C. Reichert

## Abstract

Disturbed sleep homeostatic states have been found to alter neuronal homeostasis and reduce grey matter (GM) volume. Caffeine intake that interferes with sleep homeostasis through antagonizing adenosine receptors can impair hippocampal synaptic strength, neurogenesis, as well as memory and learning in rats. In this study, reduced medial temporal GM volume was observed after daily caffeine intake in humans (3 × 150 mg × 10 days compared to 10-day placebo administration). The potential bias from reduced cerebral blood flow was controlled, and the GM reduction was independent of the change in sleep pressure. A decrease in working memory during daily caffeine intake was observed, albeit no association with the magnitude of GM changes. The findings indicate that daily caffeine intake might induce rapid cerebral plasticity and be detrimental for higher order cognitive performance in the long run. They may call into question whether the neuroprotective effects of caffeine found in acute or low dose administration in animals are generalizable onto the daily usage in humans.

## Introduction

Caffeine is the most commonly used psychostimulant worldwide and mainly consumed in forms of coffee, tea, energy drink, and soda ^1–4^. Although caffeine is mostly considered to be non-addictive, the observed physical and psychological dependence ^5,6^ consolidate its regular consumption ^7–9^ through the caffeine-induced reinforcing effects ^10^, as well as the motive to resist withdrawal symptoms ^11^ and to increase alertness ^12^. Higher alertness after acute caffeine intake ^13^ mirrors a reduced homeostatic sleep pressure, which is also evident in a reduced depth of sleep ^14^. The latter is characterized by attenuated electroencephalographic slow-wave activity (EEG SWA, 0.75 – 4.5 Hz) in non-rapid eye movement (NREM) sleep and shortened slow-wave sleep (SWS) ^14–17^.

Disturbed sleep homeostasis can not only lead to cerebral micromorphometric alterations in the mitochondria and chromatin that leads to cell death ^18–20^, but also macrostructural changes. Lower GM volumes were observed during abnormally high sleep pressure, such as during sleep deprivation or sleep fragmentation. Liu, et al. ^21^ reported reduced thalamic GM volume along with impaired cognitive performance in healthy adults after 72-hour prolonged waking compared to baseline. Dai, et al. ^22^ demonstrated a GM dynamic through the 36-hour course of sleep deprivation, where a decrease in right thalamus, right insula, right inferior parietal lobe, and bilateral somatosensory association cortex was observed by 32 hours of sleep deprivation. In clinical studies, GM volume and cortical thickness were reduced in patients with various sleep disorders (e.g. chronic insomnia ^23,24^, sleep apnea ^25^, and narcolepsy ^26^) compared to healthy controls. Furthermore, the higher symptomatic severity of insomnia and narcolepsy was commonly associated with the reduced frontal GM. The micro- and macro-morphometric changes in GM in response to sleep deprivation might reflect the disrupted adenosine-modulated cellular homeostasis, such as cardiac microtubule dynamic ^27^, astrocytic cytoskeleton arrangement ^28^, hippocampal fiber synaptic plasticity ^29^, and the robustness of cortical axons and dendrites ^30,31^.

Caffeine interferes with sleep homeostasis by dampening the accumulation of sleep pressure through the antagonism on adenosine A_1_ and A_2A_ receptors ^15,32^. Evidence in animals shows that acute or long-term caffeine consumption inhibits the long-term potentiation (LTP) ^33^, neurogenesis ^34,35^, and cell proliferation ^36^ in hippocampus, and can impair learning and memory ^34^. However, it is still not clear whether daily caffeine consumption leads to cerebral structural modifications in healthy populations through the constant impact on sleep homeostasis, despite a number of studies exploring the functional neuroprotective effect of caffeine on Parkinson’s and Alzheimer’s disease ^37,38^.

Hence, we hypothesized that, through changes in sleep homeostasis, long-term caffeine intake alters GM structures, and later, we tested exploratorily whether the GM alterations were related to changes in memory function. Cerebral GM volume was measured by magnetic resonance imaging (MRI) together with cerebral blood flow to control for its bias on MRI signals, and sleep pressure was indexed by nighttime sleep EEG slow-wave activity. Memory performance was assessed by verbal N-back tasks.

## Results

### Caffeine-induced reductions in total GM volume and in medial temporal regions

*Total* GM volume was lower in the caffeine condition compared to placebo (t_con_ = −3.59, 95% CI= [−0.009, −0.003], p < .001, **Figure 1**). A voxel-based analysis (VBA) indicated that the reduction of GM volume was most prominent in the right medial temporal lobe (*mTL*, including hippocampus; VBA cluster-level p_FWE =_ .032, **Figure 2a**; eigenvalue t_con_ = −4.35, 95% CI = [−0.012, −0.005], p < .001), along with the left frontal pole, right postcentral gyrus, right insula, and the cerebellum at trend (i.e. VBA p_FWE_ < .063, p_FDR_ = .029). No increases of GM in the caffeine compared to the placebo condition were observed.

**Figure 1.**
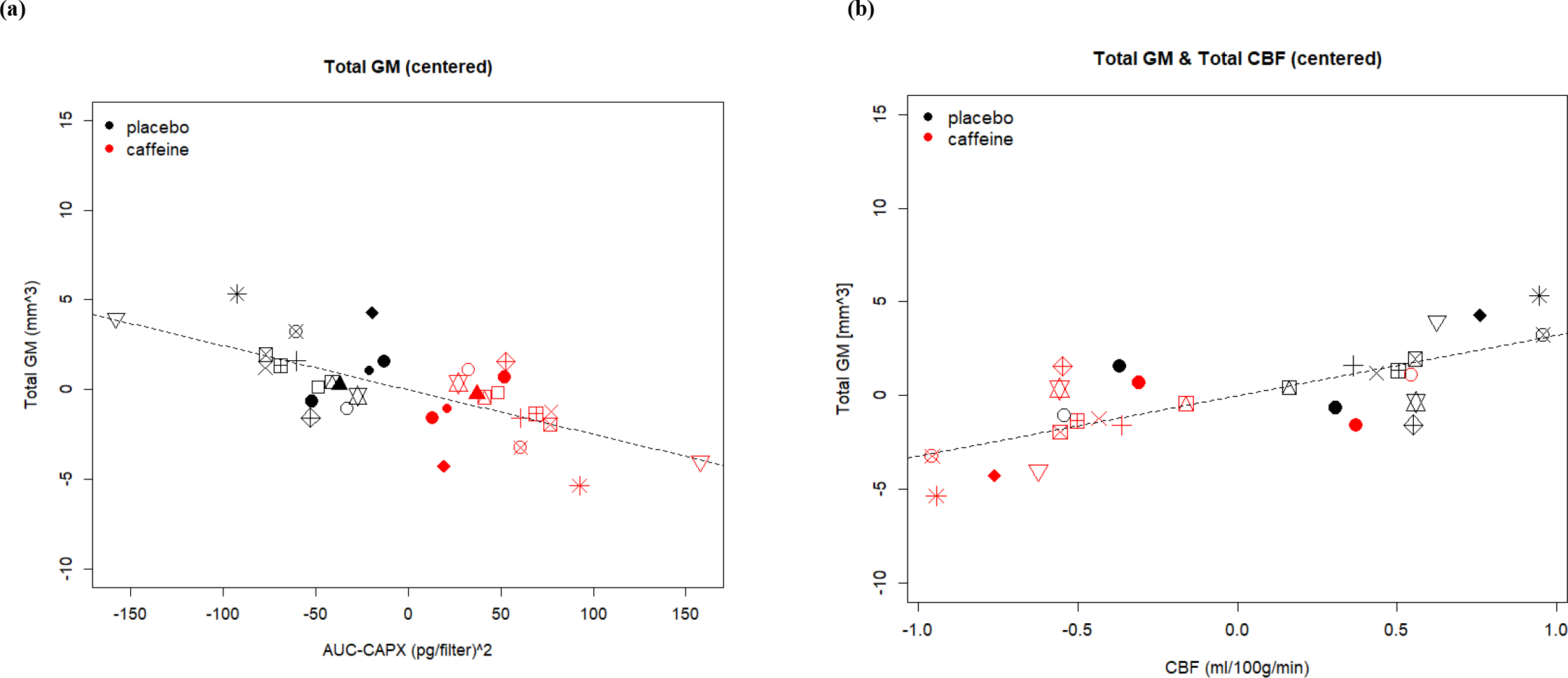
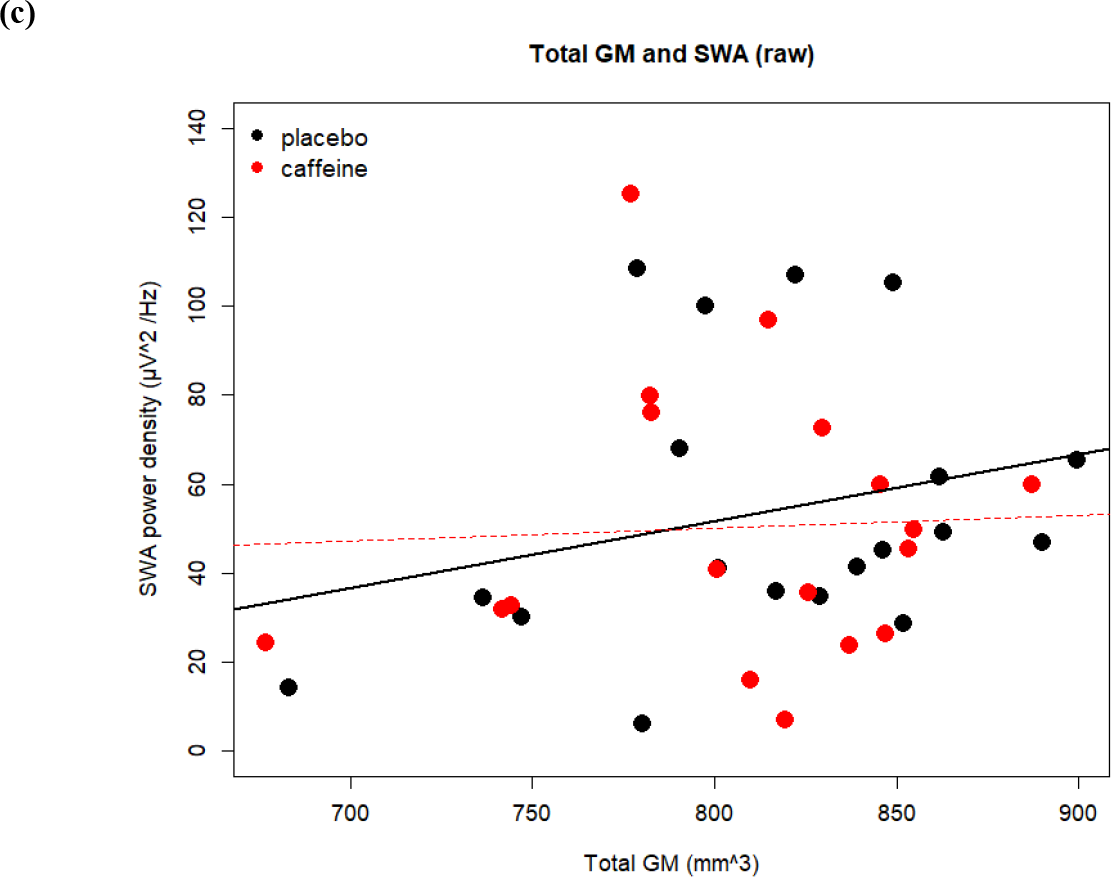
Relation of grey matter, cerebral blood flow, caffeine metabolites, and sleep slow wave activity. We use center a plot to display the changes in a specific variable to the treatment of caffeine and placebo in each subject. The values are the relative distance from the responses in each condition to the average response of a single participant, calculated as response_caff or plac_ – (response_caff_ + response_plac_)/2. Each symbol represents one participant. In two dimensions one can observe the variance of within-subject changes of two variables between conditions (by color), as well as observe the association between the changes of two variables (x and y axes). The stronger the condition effect is, the more discretely are the two-colored clouds distributed. The stronger the association between two variables is, the closer the shape is to linear. (a) and (b) *Total* GM volume is associated with AUC-CAPX negatively and with total CBF. (c) Between-subject variance of *Total* GM volume is associated with NREM SWA especially in placebo condition (presented in raw data points). Reasons for missing data points are addressed **in supplement**.

**Figure 2.**
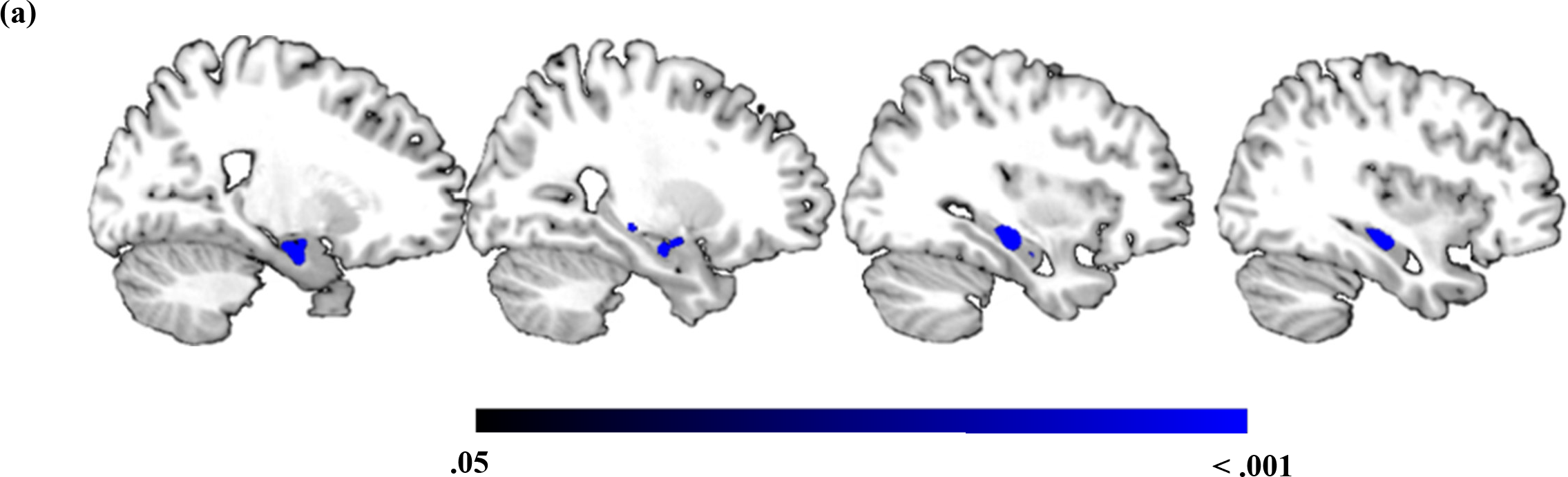
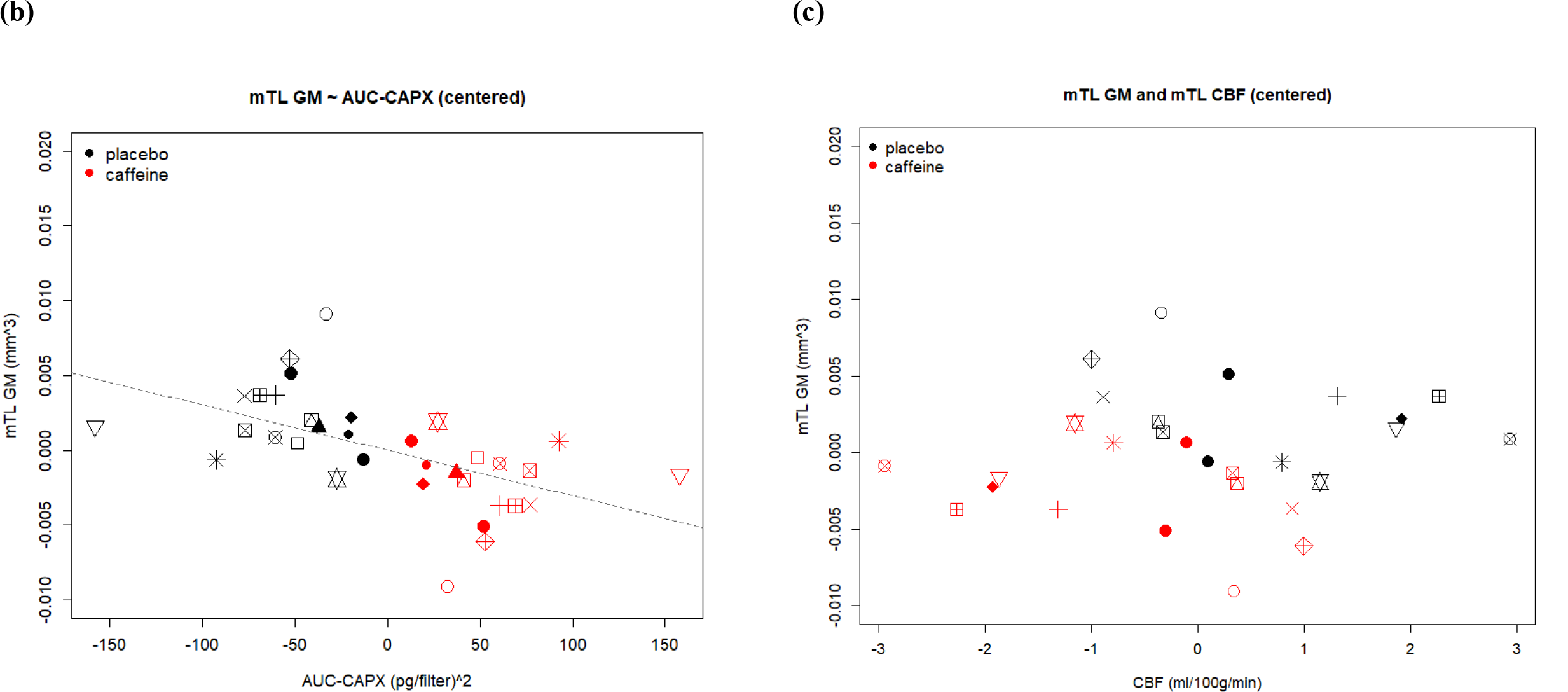
mTL GM volumetric reduction and relation with caffeine metabolites. Voxel-based and region of interest (ROI)-based approaches were both employed. The dual process allows a better spatial accuracy when first determining the condition effect on GM, while the extracted eigenvariate provides further modeling with other variables (AUC-CAPX, SWA, and CBF) to reduce bias caused by multiple comparison correction. (a) Voxel-based analysis revealed a significant GM reduction in mTL in the caffeine condition compared to placebo (color bar displayed in p_FWE_). (b) The individual responses in GM volume within the significant mTL clusters was positively associated with the levels of caffeine and paraxanthine. (c) No significant association between mTL GM and mTL CBF was found. Note that there the missing data points and reasons are addressed **in supplement**.

For further analyses, we extracted the eigenvariate of the significant cluster in the right medial temporal lobe, to which we will refer to as *mTL* GM volume. In order to support that the observed changes in *mTL* GM are induced by our treatment rather than related to a confounder, the association between the individual GM differences and the levels of caffeine and paraxanthine (AUC-CAPX) was examined (for AUC-CAPX levels during each condition across time see **supplement**). A negative association between AUC-CAPX and both *total* GM (t_AUC_ = −3.46, 95% CI = [−0.00008, −0.00002], p_AUC_ = .001; **Figure 1a**) and mTL GM (t_AUC_ = −3.83, 95% CI = [−0.004, −0.001], p_AUC_ = .001; **Figure 2b**) was found, namely the higher the AUC-CAPX the lower was the GM volume . The overview of the results and effects of coefficients are reported in **Table 1**.

**Table 1.**
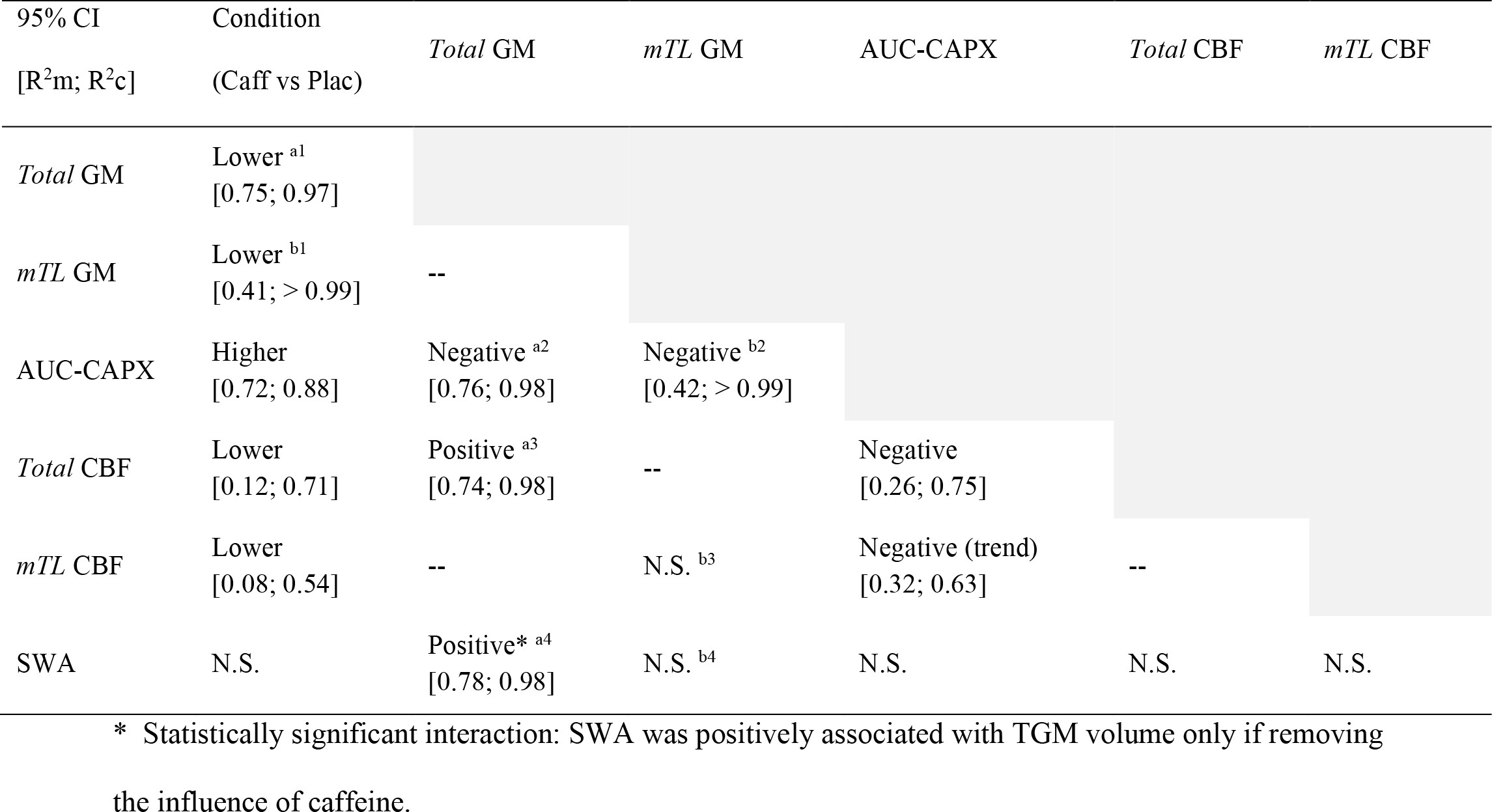
The pairwise association and its coefficient of the effects between all physiological variables. The superscripts indicate the corresponding models stated in the supplement. The coefficients of the effects (R^2^) on each model were calculated during the linear mixed model estimation, including the R^2^ marginal (R^2^m) for the coefficient of fixed effect and the R^2^ condition (R^2^c) for the coefficient for fixed + random effects.

### Caffeine-induced differences in the association of SWA and GM volumetric reductions

In a next step, it was examined whether SWA-indexed sleep pressure is reduced during long-term caffeine intake compared to placebo, and whether the changes in GM were mediated by SWA. Since SWA in the first NREM episode shows a high sensitivity to acute caffeine intake ^17,39^, this specific segment of the night was adopted to index homeostatic sleep-pressure in our study. The overview of the results and effects of coefficients are reported in **Table 1**.

No significant change in NREM SWA after 9 days of daily caffeine intake compared to 9 days of daily placebo intake was observed (t_con_ = −0.87, p_con_ = 0.386, **see supplement**). Yet, overall SWA was positively associated with *total* GM (t_SWA_ = 2.88, 95% CI = [0.00008, 0.0004], p_SWA_ = .004, **Figure 1c**), and a significant interaction between condition and SWA indicated a stronger association in the placebo condition (t_inter_ = −4.709, 95% CI = [−0.0002, −0.0001], p_inter_ < .001). In the regional *mTL* GM differences, no associations between SWA and GM volumetric reductions were seen (t_SWA_ = 0.09, p_SWA_ = .931).

### CBF response during daily caffeine consumption

When estimating GM tissue volume, caffeine-induced reductions in regional CBF can regionally bias the MRI signal distribution and therefore the tissue probability maps ^40–42^. Thus, we first examined the CBF response during caffeine intake compared to placebo, followed by the inspection of its contribution to the observed GM changes. The overview of the results and effects of coefficients are reported in **Table 1**.

A significant reduction in total CBF was found in the caffeine compared to the placebo condition (t_con_ = −5.2, 95% CI = [−0.119, −0.054], p_con_ < .001). Voxel-wise analysis revealed that reductions occurred mainly in the midline, including cuneus, precuneus, and subcortical regions (p_FWE_all_ < .05, **Figure 3**). Importantly, also in the region where a caffeine-associated GM volumetric reduction was observed, CBF was also lower in the caffeine condition compared to placebo (t_con_ = −2.17, 95% CI = [−2.644, −0.132], p_con_ = .048). AUC-CAPX was negatively associated with both total CBF (t_AUC_ = −6.33, 95% CI = [−0.0009, −0.0005], p_AUC =_ .001) and mTL CBF (t_AUC_ = −2.77, 95% CI = [−0.021, −0.004], p_AUC_ = .055).

**Figure 3.**
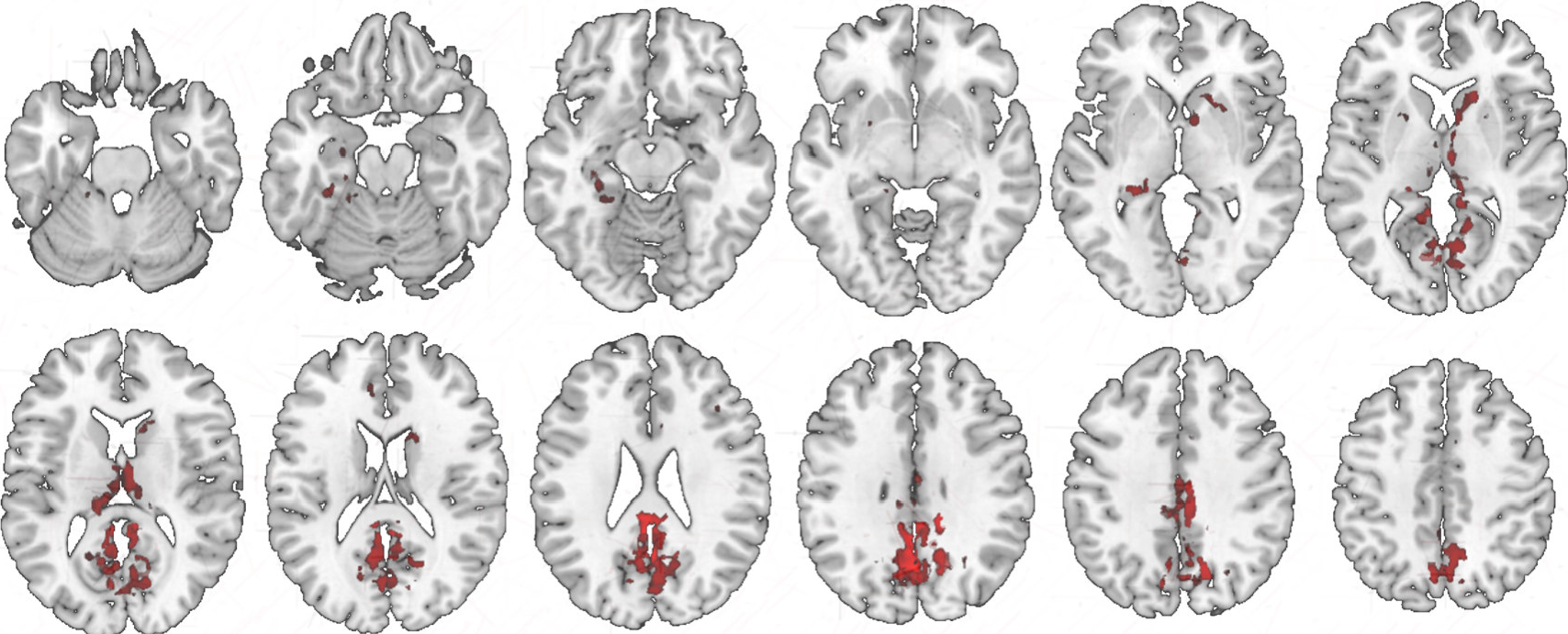
Caffeine-associated reduction in CBF. Regions (in red) showing a reduction in CBF after caffeine intake. Most prominent reduction were observed in cuneus, precuneus, and subcortical regions (p_FWE_ < .05).

Despite a reduction in both total and *mTL* CBF, the associations with GM volume were different. A positive association between the reduction of *total* CBF and *total* GM volume was observed (t_CBF_ = 4.70, 95% CI = [0.004, 0.009], p_CBF_ < .001). Moreover, the variance of CBF mediated caffeine-associated reductions on *total* GM: Including *total* CBF as a covariate fully accounted for the main effect of condition on *total* GM estimation (t_CBF_ = 2.82, p_CBF_ = .005; t_con_ = −0.77, p_con_ = .441, **Figure 1b**). In contrast, CBF did not account for the caffeine-induced volumetric reductions in *mTL* GM, as indicated by multi-modal VBA with linear mixed model. Consistently, in the ROI-based analysis, no association between the caffeine-associated changes in *mTL* CBF and in *mTL* GM was found (t_CBF_ = −1.67, p_CBF_ = .122, **Figure 2c**). Thus, reasonably, no mediation was exerted by *mTL* CBF on the *mTL* GM volumetric reduction (t_CBF_ = −1.88, p_CBF_ = .083; t_con =_ −4.80, p_con_ < .001).

### Working memory performance during daily caffeine intake

In a last step, a possible functional implication of the caffeine-induced *mTL* GM reductions was explored. To do so, performance in a verbal n-back task with two levels of difficulty (0-back, referred to as low load; 3-back, referred to as high load) was analyzed with adjustment for the order of the conditions. The task was administered every 4 hours during 12.5 hours before and during the MRI measurements. The differences in average accuracy and reaction time (RT) of four attempts between the caffeine and placebo condition were examined. The overview of the results, 95% CI, and effects of coefficients are reported in **the supplement**.

Compared to placebo, lower accuracy was found in both 0-back (t_con_ = −2.33, p_con_ = .020, **Figure 4**) and 3-back (t_con_ = −2.02, p_con_ = .044, **Figure 4**) performance in the caffeine condition, as well as a lower net accuracy of 3-back, i.e. corrected for the baseline response of 0-back performance (t_con_ = −1.97, 95% CI = [−0.0355, −0.00010], p_con_ = .049; R^2^m = .41, R^2^c = .85). The reductions in accuracy were not linked to the reductions in total or regional GM in caffeine compared to placebo. Moreover, in contrast to the worse accuracy in the caffeine condition compared to placebo, a higher AUC-CAPX level within the caffeine condition was positively associated with a better net 3-back accuracy (t_AUC_ = 3.58, p_AUC_ < .001). No significant association of AUC-CAPX and 0-back accuracy was observed. On the other hand, while there was no difference found in the net overall RT and the RT of hits and false alarm responses between two conditions (t_all_ < 1.88, p_all_ > .60), the RT of missed (t_con_ = 2.75, p_con_ = .006) and correction rejections (t_con =_1.95, p_con_ = .051) were longer in caffeine condition compared to placebo. The net RT in all types of responses, however, was negatively associated with AUC-CAPX (t_all_ < −2.44, p_all_ < .015).

**Figure 4.**
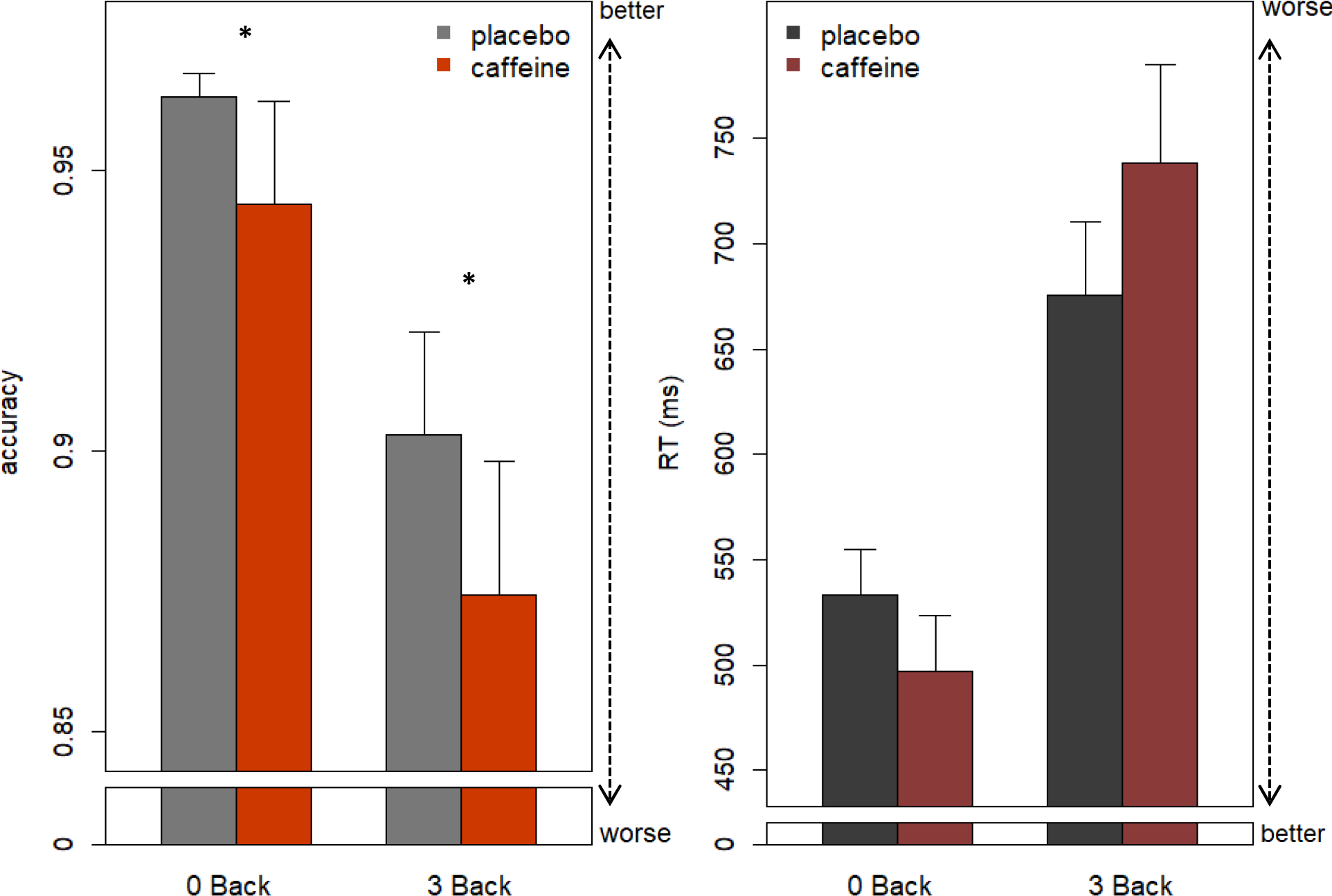
Differences in verbal working memory accuracy and RT between caffeine and placebo. In the caffeine condition, the accuracy was significantly worse in both 0-back and 3-back compared to placebo. We observed no significant difference in overall RTs.

## Discussion

Our study examined whether daily caffeine intake alters cerebral morphometry, and whether these changes are mediated by homeostatic sleep pressure, as indexed by sleep EEG slow-wave activity (SWA). We observed reductions in total grey matter (GM) volume, which were accounted for by cerebral blood flow (CBF), suggesting a strong necessity to strictly control for both caffeine intake and CBF in future studies of GM volume. However, we also observed caffeine-induced concentration-dependent reductions in GM volume in a cluster within the medial temporal lobe (including hippocampus, parahippocampus, fusiform gyrus), which were not accounted for by the reduction in CBF. In contrast to our hypothesis, daily caffeine intake did not reduce EEG slow-wave activity during the night sleep, nor were the GM reductions directly mediated by alterations of sleep SWA. Caffeine might, thus, exerts an effect on cerebral plasticity through a parallel pathway to its influence on sleep, as discussed in the following sections. In a nutshell, our data indicate that daily intake of caffeine, as observed in around 80% of the population in the world ^43^, challenges GM plasticity. Together with the observed impairments of working memory performance, they strongly question earlier-reported benefits of daily caffeine intake especially in healthy populations.

A caffeine-induced lower GM volume may reflect multiple possible responses, one main assumption is neuronal damage. It has been observed that daily or high-dose caffeine intake weakens A_2A_R affinity for caffeine ^44^ and lead to an upregulated A_2A_R availability ^45,46^, consequently maintains boosting in glutamate signaling and prolonged post firing of hippocampal pyramidal cells ^44,47,48^. Furthermore, the strengthened A_2A_R agonism decreases the affinity of A_1_R for caffeine on the A_1_-A_2A_ receptor heteromers, which is speculated to result in a tolerance effect in behaviors after daily use of caffeine ^44,49^ and, potentially as observed in our study, SWA.

Sleep SWA is positively linked with adenosine binding and dissipation during sleep ^50^. SWA renormalizes the saturated synaptic capacity and recover the brain neurons from the energy consumption during prior wakefulness, suggested by the synaptic homeostasis hypothesis (SHY) ^51^. The strength of synaptic potentiation can be regained only if the renormalization of synaptic strength has occurred during sleep ^52^. Our data suggest that chronic caffeine intake might affect this homeostatic balance. We observed a between-subject concentration-dependent enhancement on memory performance that might reflect the boosting effect of caffeine on cerebral activities. At the same time, however, there was lack of a commensurate response in SWA as the due synaptic recovery from the toned-up neuronal strength. A deficient synaptic restoration followed by the continuous administration of caffeine on the next day may presumably maintain the cellular stress, inhibit the functions of hippocampal pyramidal cells ^33–35^ due to exhaustion, and, in the long run, impair memory performance as observed in our study as well as other animal studies on daily administration of caffeine ^34^ and adenosine antagonists ^53^.

Beside a change in neurons, the lower GM observed in our study can also result from changes in non-neuronal cells and/or in cerebral vasculature ^54^. In oncological studies, caffeine has been applied to induce apoptosis of glial cells ^55^. Despite an absence of direct evidence on the effect of daily caffeine intake, the effects of adenosine and its A_2A_ receptors on the release of growth factors have been strongly suggested ^38^, which in turn modulate the proliferation of astrocytes and can influence angiogenesis ^56^.

Changes in GM volume, as derived by VBM, can also be simply confounded by differences in CBF ^40–42^. Accordingly, in the present study, CBF accounted for *total* GM changes induced by caffeine intake. These apparent *total GM* changes emphasize the importance of controlling for caffeine consumption in repeated fMRI measures, especially with a region of interests in the most impacted regions (cuneus, precuneus, cerebellum, and subcortices). The observation of the typical caffeine-induced reductions in CBF ^42,57,58^ in our study, moreover add to the current knowledge that the history of caffeine intake at least within 4.5 hours strongly impacts on both CBF and apparent tissue change.

Our treatment did not elicit a significant effect on SWA in the first NREM sleep episode. The contradiction to earlier findings^14, 16,17^ can be reconciled within two assumptions: a timing-dependent influence of caffeine on sleep, and a potential development of tolerance in response to the repeated daily intake. Earlier studies indicate that caffeine exerts strong reduction in NREM SWA when caffeine was administered close to bedtime ^16,17^ while the effects were considerably weaker after morning administration ^59^. Moreover, a tolerance effect to caffeine treatment was found in other sleep features during a 11-day course ^60^, which might similarly apply to SWA.

The combination of presence and absence of caffeine effects on CBF and SWA respectively further suggest distinct responses of A_1_R and A_2A_R in developing tolerance to daily intake ^46,61–64^. While the antagonism on A_2A_ and A_2B_ receptors plays a predominant role on vasoconstriction ^65,66^, the sleep homeostatic regulation is primarily modulated by A_1_R through the cholinergic signaling in basal forebrain ^14,15,67^ together with an indirect modulation from A_2A_R ^68^. Therefore, while daily intake of caffeine reduces CBF through the blockade of A_2A_R without developing complete tolerance, the tolerance response of A_1_R might result in the divergent response in SWA. Furthermore, chronic caffeine treatment is found to strengthen A_2A_R agonism on its A_1_R-heteromers, which leads to a decrease in caffeine binding to A_1_R and speculatively result in a tolerance effect in behaviors ^44,49^, and potentially, SWA.

Notably, we also observed a positive between-subject association between SWA and *total* GM volume particularly in the placebo condition, which might reflect the individual variance of brain maturity during young adulthood ^69^. However, the attenuated association in caffeine condition might simply be caused by the increased variance due to the bias of CBF after caffeine intake. On the contrary, GM in the mTL region sensitive to our treatment did not exhibit this association, which potentially reflects region-specific mechanisms of plasticity during sleep ^70^.

As the first study about caffeine effects on GM in young healthy adults, our results may seem contradictory to earlier evidence on its neuroprotective effect on cognitive impairment ^71,72^. However, studies that demonstrated the neuroprotective effects of caffeine/adenosine antagonists were mostly based on preventive models, where one observed the efficacy of the agent in reversing neurodegeneration or inflammation induced by kinase, transgenicity, or trauma ^8,73,74^. The effects therefore cannot be generalized and thus have left the puzzles on how the constant interference of caffeine on a healthy adenosinergic system would alter the intrinsic neurotransmission. Moreover, there is a considerable inconsistency in the observed neuroprotective effects of caffeine and other adenosine antagonists^75^. For instance, a high-dose administration of A_2A_R antagonists can induce excitotoxicity acutely *in vivo* and *ex vivo* ^76,77^. A peripheral administration and a direct neuronal injection of identical selective A_2A_ antagonist might also lead to opposite outcomes on the neuroprotective effects ^78,79^. A recent study suggested that age-related cognitive and neuronal declines were ameliorated by chronic caffeine consumption, yet also with lower dose (5 mg/kg/day) as compared to our study ^80^. Given the distinct responses and interaction between the subtypes of adenosine receptors, the difference between pre- and post-synaptic administration, and the dose-dependent effect, it is of importance to carefully compare the study designs that focused on a similar research question.

Our study also bears some limitations that require a careful interpretation. Firstly, although a power calculation indicated a sufficient sample size, we came across data loss due to technical reasons, which lowers the signal-to-noise ratio to detect a relatively small effect. To address this issue, a non-parametric permutation test was therefore employed in all the voxel-based analyses as an initial step, and the subsequent analyses were established upon its results. Secondly, one might argue that the absence of a significant difference in SWA might be due to a genetically predisposed insensitivity to caffeine of the selected population ^81^. However, as the withdrawal from daily intake of 450 mg of caffeine has been shown to induce clear-cut responses in the sleep-homeostatic regulation of these participants ^82^, it seems unlikely that the selected population was insensitive to the stimulant. Notably, the habitual amount was calculated from all types of caffeinated diets, not only coffee intake but also tea, chocolate drinks, energy drinks, soda, and so on.

Overall, our findings derived from this first within-subject laboratory study yield an insight on the mTL GM plasticity induced by the repeated intake of caffeine in a long-term course. Reductions in GM volume and the cooccurring impairment in working memory performance underline the importance of carefully questioning the consequences of habitual daily intake of a psychostimulant, freely available all over the world.

## Acknowledgement

We sincerely appreciate the contribution of our interns Andrea Schumacher, Laura Tincknell, M.Sc. student Sven Leach, and all the study helpers assisting in the experiment. We also thank Dr. med. Corrado Garbazza and Dr. med. Helen Slawik for the health check during screening process. We acknowledge gratefully the collaboration with Prof. Christopher Gerner and Dr. Samuel Meier at University of Vienna, together with the assistance of M.Sc. Max Feuerstein, and Dr. Rupert Mayer for the measurement of fingertip sweats. We especially appreciate all our participants for their volunteering and cooperation. This project is financed by *Swiss National Science Foundation (320030-163058).*

## References

1 Mitchell, D. C., Knight, C. A., Hockenberry, J., Teplansky, R. & Hartman, T. J. Beverage caffeine intakes in the U.S. Food and chemical toxicology : an international journal published for the British Industrial Biological Research Association 63, 136–142, doi:10.1016/j.fct.2013.10.042 (2014).

2 Frary, C. D., Johnson, R. K. & Wang, M. Q. Food sources and intakes of caffeine in the diets of persons in the United States. Journal of the American Dietetic Association 105, 110–113, doi:10.1016/j.jada.2004.10.027 (2005).

3 Reyes, C. M. & Cornelis, M. C. Caffeine in the Diet: Country-Level Consumption and Guidelines. Nutrients 10, doi:10.3390/nu10111772 (2018).

4 Barone, J. J. & Roberts, H. R. Caffeine consumption. Food and chemical toxicology : an international journal published for the British Industrial Biological Research Association 34, 119–129 (1996).

5 Nehlig, A. Are we dependent upon coffee and caffeine? A review on human and animal data. Neuroscience and biobehavioral reviews 23, 563–576 (1999).

6 Mills, L., Boakes, R. A. & Colagiuri, B. Placebo caffeine reduces withdrawal in abstinent coffee drinkers. Journal of psychopharmacology (Oxford, England) 30, 388–394, doi:10.1177/0269881116632374 (2016).

7 Fredholm, B. B., Battig, K., Holmen, J., Nehlig, A. & Zvartau, E. E. Actions of caffeine in the brain with special reference to factors that contribute to its widespread use. Pharmacological reviews 51, 83–133 (1999).

8 Ferre, S. An update on the mechanisms of the psychostimulant effects of caffeine. Journal of neurochemistry 105, 1067–1079, doi:10.1111/j.1471-4159.2007.05196.x (2008).

9 Ferre, S. Mechanisms of the psychostimulant effects of caffeine: implications for substance use disorders. Psychopharmacology 233, 1963–1979, doi:10.1007/s00213-016-4212-2 (2016).

10 Griffiths, R. R. & Woodson, P. P. Reinforcing effects of caffeine in humans. The Journal of pharmacology and experimental therapeutics 246, 21–29 (1988).

11 Juliano, L. M. & Griffiths, R. R. A critical review of caffeine withdrawal: empirical validation of symptoms and signs, incidence, severity, and associated features. Psychopharmacology (Berl) 176, 1–29, doi:10.1007/s00213-004-2000-x (2004).

12 Mahoney, C. R. et al. Intake of caffeine from all sources and reasons for use by college students. Clin Nutr, doi:10.1016/j.clnu.2018.04.004 (2018).

13 Einother, S. J. & Giesbrecht, T. Caffeine as an attention enhancer: reviewing existing assumptions. Psychopharmacology 225, 251–274, doi:10.1007/s00213-012-2917-4 (2013).

14 Clark, I. & Landolt, H. P. Coffee, caffeine, and sleep: A systematic review of epidemiological studies and randomized controlled trials. Sleep medicine reviews 31, 70–78, doi:10.1016/j.smrv.2016.01.006 (2017).

15 Urry, E. & Landolt, H. P. Adenosine, caffeine, and performance: from cognitive neuroscience of sleep to sleep pharmacogenetics. Current topics in behavioral neurosciences 25, 331–366, doi:10.1007/7854_2014_274 (2015).

16 Drapeau, C. et al. Challenging sleep in aging: the effects of 200 mg of caffeine during the evening in young and middle-aged moderate caffeine consumers. Journal of sleep research 15, 133–141, doi:10.1111/j.1365-2869.2006.00518.x (2006).

17 Landolt, H. P., Dijk, D. J., Gaus, S. E. & Borbely, A. A. Caffeine reduces low-frequency delta activity in the human sleep EEG. Neuropsychopharmacology : official publication of the American College of Neuropsychopharmacology 12, 229–238, doi:10.1016/0893-133x(94)00079-f (1995).

18 Zhao, Z., Zhao, X. & Veasey, S. C. Neural Consequences of Chronic Short Sleep: Reversible or Lasting? Frontiers in neurology 8, 235, doi:10.3389/fneur.2017.00235 (2017).

19 Abushov, B. M. Morphofunctional analysis of the effects of total sleep deprivation on the CNS in rats. Neuroscience and behavioral physiology 40, 403–409, doi:10.1007/s11055-010-9271-y (2010).

20 Zhao, H. et al. Frontal cortical mitochondrial dysfunction and mitochondria-related beta-amyloid accumulation by chronic sleep restriction in mice. Neuroreport 27, 916–922, doi:10.1097/wnr.0000000000000631 (2016).

21 Liu, C., Kong, X. Z., Liu, X., Zhou, R. & Wu, B. Long-term total sleep deprivation reduces thalamic gray matter volume in healthy men. Neuroreport 25, 320–323, doi:10.1097/wnr.0000000000000091 (2014).

22 Dai, X. J. et al. Plasticity and Susceptibility of Brain Morphometry Alterations to Insufficient Sleep. Frontiers in psychiatry 9, 266, doi:10.3389/fpsyt.2018.00266 (2018).

23 Altena, E., Vrenken, H., Van Der Werf, Y. D., van den Heuvel, O. A. & Van Someren, E. J. Reduced orbitofrontal and parietal gray matter in chronic insomnia: a voxel-based morphometric study. Biological psychiatry 67, 182–185, doi:10.1016/j.biopsych.2009.08.003 (2010).

24 Joo, E. Y. et al. Brain Gray Matter Deficits in Patients with Chronic Primary Insomnia. Sleep 36, 999–1007, doi:10.5665/sleep.2796 (2013).

25 Baril, A. A. et al. Gray Matter Hypertrophy and Thickening with Obstructive Sleep Apnea in Middle-aged and Older Adults. American journal of respiratory and critical care medicine 195, 1509–1518, doi:10.1164/rccm.201606-1271OC (2017).

26 Joo, E. Y. et al. Analysis of cortical thickness in narcolepsy patients with cataplexy. Sleep 34, 1357–1364, doi:10.5665/sleep.1278 (2011).

27 Fassett, J. T. et al. Adenosine regulation of microtubule dynamics in cardiac hypertrophy. Am J Physiol Heart Circ Physiol 297, H523–532, doi:10.1152/ajpheart.00462.2009 (2009).

28 Abbracchio, M. P. et al. The A3 adenosine receptor induces cytoskeleton rearrangement in human astrocytoma cells via a specific action on Rho proteins. Annals of the New York Academy of Sciences 939, 63–73 (2001).

29 Kukley, M., Schwan, M., Fredholm, B. B. & Dietrich, D. The role of extracellular adenosine in regulating mossy fiber synaptic plasticity. The Journal of neuroscience : the official journal of the Society for Neuroscience 25, 2832–2837, doi:10.1523/jneurosci.4260-04.2005 (2005).

30 Ribeiro, F. F. & Sebastiao, A. M. Adenosine A2A receptors in neuronal outgrowth: a target for nerve regeneration? Neural regeneration research 11, 706–708, doi:10.4103/1673-5374.182683 (2016).

31 Ribeiro, F. F. et al. Axonal elongation and dendritic branching is enhanced by adenosine A2A receptors activation in cerebral cortical neurons. Brain structure & function 221, 2777–2799, doi:10.1007/s00429-015-1072-1 (2016).

32 Elmenhorst, D., Meyer, P. T., Matusch, A., Winz, O. H. & Bauer, A. Caffeine occupancy of human cerebral A1 adenosine receptors: in vivo quantification with 18F-CPFPX and PET. Journal of nuclear medicine : official publication, Society of Nuclear Medicine 53, 1723–1729, doi:10.2967/jnumed.112.105114 (2012).

33 Blaise, J. H., Park, J. E., Bellas, N. J., Gitchell, T. M. & Phan, V. Caffeine consumption disrupts hippocampal long-term potentiation in freely behaving rats. Physiological reports 6, doi:10.14814/phy2.13632 (2018).

34 Han, M. E. et al. Inhibitory effects of caffeine on hippocampal neurogenesis and function. Biochemical and biophysical research communications 356, 976–980, doi:10.1016/j.bbrc.2007.03.086 (2007).

35 Wentz, C. T. & Magavi, S. S. Caffeine alters proliferation of neuronal precursors in the adult hippocampus. Neuropharmacology 56, 994–1000, doi:10.1016/j.neuropharm.2009.02.002 (2009).

36 Kochman, L. J., Fornal, C. A. & Jacobs, B. L. Suppression of hippocampal cell proliferation by short-term stimulant drug administration in adult rats. The European journal of neuroscience 29, 2157–2165, doi:10.1111/j.1460-9568.2009.06759.x (2009).

37 Chen, J. F. Adenosine receptor control of cognition in normal and disease. International review of neurobiology 119, 257–307, doi:10.1016/b978-0-12-801022-8.00012-x (2014).

38 Cunha, R. A. How does adenosine control neuronal dysfunction and neurodegeneration? Journal of neurochemistry 139, 1019–1055, doi:10.1111/jnc.13724 (2016).

39 Carrier, J. et al. Effects of caffeine on daytime recovery sleep: A double challenge to the sleep-wake cycle in aging. Sleep medicine 10, 1016–1024, doi:10.1016/j.sleep.2009.01.001 (2009).

40 Laurienti, P. J. et al. Relationship between caffeine-induced changes in resting cerebral perfusion and blood oxygenation level-dependent signal. AJNR. American journal of neuroradiology 24, 1607–1611 (2003).

41 Ge, Q. et al. Short-term apparent brain tissue changes are contributed by cerebral blood flow alterations. PloS one 12, e0182182, doi:10.1371/journal.pone.0182182 (2017).

42 Field, A. S., Laurienti, P. J., Yen, Y. F., Burdette, J. H. & Moody, D. M. Dietary caffeine consumption and withdrawal: confounding variables in quantitative cerebral perfusion studies? Radiology 227, 129–135, doi:10.1148/radiol.2271012173 (2003).

43 James, J. Understanding Caffeine: a Biobehavioral Analysis. (Sage Publications, 1997).

44 Ciruela, F. et al. Presynaptic control of striatal glutamatergic neurotransmission by adenosine A1-A2A receptor heteromers. The Journal of neuroscience : the official journal of the Society for Neuroscience 26, 2080–2087, doi:10.1523/jneurosci.3574-05.2006 (2006).

45 Varani, K. et al. Caffeine intake induces an alteration in human neutrophil A2A adenosine receptors. Cellular and molecular life sciences : CMLS 62, 2350–2358, doi:10.1007/s00018-005-5312-z (2005).

46 Johansson, B., Georgiev, V., Lindstrom, K. & Fredholm, B. B. A1 and A2A adenosine receptors and A1 mRNA in mouse brain: effect of long-term caffeine treatment. Brain research 762, 153–164 (1997).

47 Fontinha, B. M., Diogenes, M. J., Ribeiro, J. A. & Sebastiao, A. M. Enhancement of long-term potentiation by brain-derived neurotrophic factor requires adenosine A2A receptor activation by endogenous adenosine. Neuropharmacology 54, 924–933, doi:10.1016/j.neuropharm.2008.01.011 (2008).

48 Rombo, D. M. et al. Synaptic mechanisms of adenosine A2A receptor-mediated hyperexcitability in the hippocampus. Hippocampus 25, 566–580, doi:10.1002/hipo.22392 (2015).

49 Ferre, S. et al. Adenosine A1-A2A receptor heteromers: new targets for caffeine in the brain. Frontiers in bioscience : a journal and virtual library 13, 2391–2399 (2008).

50 Greene, R. W., Bjorness, T. E. & Suzuki, A. The adenosine-mediated, neuronal-glial, homeostatic sleep response. Current opinion in neurobiology 44, 236–242, doi:10.1016/j.conb.2017.05.015 (2017).

51 Tononi, G. & Cirelli, C. Sleep and synaptic homeostasis: a hypothesis. Brain research bulletin 62, 143–150 (2003).

52 Tononi, G. & Cirelli, C. Sleep and the price of plasticity: from synaptic and cellular homeostasis to memory consolidation and integration. Neuron 81, 12–34, doi:10.1016/j.neuron.2013.12.025 (2014).

53 Von Lubitz, D. K., Paul, I. A., Bartus, R. T. & Jacobson, K. A. Effects of chronic administration of adenosine A1 receptor agonist and antagonist on spatial learning and memory. European journal of pharmacology 249, 271–280, doi:10.1016/0014-2999(93)90522-j (1993).

54 Zatorre, R. J., Fields, R. D. & Johansen-Berg, H. Plasticity in gray and white: neuroimaging changes in brain structure during learning. Nat Neurosci 15, 528–536, doi:10.1038/nn.3045 (2012).

55 Li, Y. et al. Autophagy mediated by endoplasmic reticulum stress enhances the caffeine-induced apoptosis of hepatic stellate cells. International journal of molecular medicine 40, 1405–1414, doi:10.3892/ijmm.2017.3145 (2017).

56 Fredholm, B. B. Adenosine, an endogenous distress signal, modulates tissue damage and repair. Cell death and differentiation 14, 1315–1323, doi:10.1038/sj.cdd.4402132 (2007).

57 Vidyasagar, R., Greyling, A., Draijer, R., Corfield, D. R. & Parkes, L. M. The effect of black tea and caffeine on regional cerebral blood flow measured with arterial spin labeling. Journal of cerebral blood flow and metabolism : official journal of the International Society of Cerebral Blood Flow and Metabolism 33, 963–968, doi:10.1038/jcbfm.2013.40 (2013).

58 Merola, A. et al. Mapping the pharmacological modulation of brain oxygen metabolism: The effects of caffeine on absolute CMRO2 measured using dual calibrated fMRI. Neuroimage 155, 331–343, doi:10.1016/j.neuroimage.2017.03.028 (2017).

59 Landolt, H. P., Werth, E., Borbely, A. A. & Dijk, D. J. Caffeine intake (200 mg) in the morning affects human sleep and EEG power spectra at night. Brain research 675, 67–74 (1995).

60 Bonnet, M. H. & Arand, D. L. Caffeine use as a model of acute and chronic insomnia. Sleep 15, 526–536 (1992).

61 Popoli, P., Reggio, R. & Pezzola, A. Effects of SCH 58261, an adenosine A(2A) receptor antagonist, on quinpirole-induced turning in 6-hydroxydopamine-lesioned rats. Lack of tolerance after chronic caffeine intake. Neuropsychopharmacology : official publication of the American College of Neuropsychopharmacology 22, 522–529, doi:10.1016/s0893-133x(99)00144-x (2000).

62 Halldner, L., Lozza, G., Lindstrom, K. & Fredholm, B. B. Lack of tolerance to motor stimulant effects of a selective adenosine A(2A) receptor antagonist. European journal of pharmacology 406, 345–354 (2000).

63 Karcz-Kubicha, M. et al. Involvement of adenosine A1 and A2A receptors in the motor effects of caffeine after its acute and chronic administration. Neuropsychopharmacology : official publication of the American College of Neuropsychopharmacology 28, 1281–1291, doi:10.1038/sj.npp.1300167 (2003).

64 Johansson, B. et al. Effect of long term caffeine treatment on A1 and A2 adenosine receptor binding and on mRNA levels in rat brain. Naunyn-Schmiedeberg’s archives of pharmacology 347, 407–414 (1993).

65 Ngai, A. C., Coyne, E. F., Meno, J. R., West, G. A. & Winn, H. R. Receptor subtypes mediating adenosine-induced dilation of cerebral arterioles. Am J Physiol Heart Circ Physiol 280, H2329–2335, doi:10.1152/ajpheart.2001.280.5.H2329 (2001).

66 Meno, J. R., Crum, A. V. & Winn, H. R. Effect of adenosine receptor blockade on pial arteriolar dilation during sciatic nerve stimulation. Am J Physiol Heart Circ Physiol 281, H2018–2027, doi:10.1152/ajpheart.2001.281.5.H2018 (2001).

67 Pelligrino, D. A., Xu, H. L. & Vetri, F. Caffeine and the control of cerebral hemodynamics. Journal of Alzheimer’s disease : JAD 20 Suppl 1, S51–62, doi:10.3233/jad-2010-091261 (2010).

68 Ferre, S. Role of the central ascending neurotransmitter systems in the psychostimulant effects of caffeine. Journal of Alzheimer’s disease : JAD 20 Suppl 1, S35–49, doi:10.3233/jad-2010-1400 (2010).

69 Buchmann, A. et al. EEG sleep slow-wave activity as a mirror of cortical maturation. Cerebral cortex (New York, N.Y. : 1991) 21, 607–615, doi:10.1093/cercor/bhq129 (2011).

70 Niethard, N., Burgalossi, A. & Born, J. Plasticity during Sleep Is Linked to Specific Regulation of Cortical Circuit Activity. Frontiers in neural circuits 11, 65, doi:10.3389/fncir.2017.00065 (2017).

71 Arendash, G. W. et al. Caffeine protects Alzheimer’s mice against cognitive impairment and reduces brain beta-amyloid production. Neuroscience 142, 941–952, doi:10.1016/j.neuroscience.2006.07.021 (2006).

72 Arendash, G. W. et al. Caffeine reverses cognitive impairment and decreases brain amyloid-beta levels in aged Alzheimer’s disease mice. Journal of Alzheimer’s disease : JAD 17, 661–680, doi:10.3233/jad-2009-1087 (2009).

73 Cunha, R. A. Adenosine as a neuromodulator and as a homeostatic regulator in the nervous system: different roles, different sources and different receptors. Neurochemistry international 38, 107–125 (2001).

74 Rivera-Oliver, M. & Diaz-Rios, M. Using caffeine and other adenosine receptor antagonists and agonists as therapeutic tools against neurodegenerative diseases: a review. Life sciences 101, 1–9, doi:10.1016/j.lfs.2014.01.083 (2014).

75 Dai, S. S. & Zhou, Y. G. Adenosine 2A receptor: a crucial neuromodulator with bidirectional effect in neuroinflammation and brain injury. Reviews in the neurosciences 22, 231–239, doi:10.1515/rns.2011.020 (2011).

76 Popoli, P. et al. Blockade of striatal adenosine A2A receptor reduces, through a presynaptic mechanism, quinolinic acid-induced excitotoxicity: possible relevance to neuroprotective interventions in neurodegenerative diseases of the striatum. The Journal of neuroscience : the official journal of the Society for Neuroscience 22, 1967–1975 (2002).

77 Tebano, M. T. et al. Adenosine A2A receptor blockade differentially influences excitotoxic mechanisms at pre- and postsynaptic sites in the rat striatum. Journal of neuroscience research 77, 100–107, doi:10.1002/jnr.20138 (2004).

78 Jones, P. A., Smith, R. A. & Stone, T. W. Protection against hippocampal kainate excitotoxicity by intracerebral administration of an adenosine A2A receptor antagonist. Brain research 800, 328–335 (1998).

79 Jones, P. A., Smith, R. A. & Stone, T. W. Protection against kainate-induced excitotoxicity by adenosine A2A receptor agonists and antagonists. Neuroscience 85, 229–237 (1998).

80 Vila-Luna, S. et al. Chronic caffeine consumption prevents cognitive decline from young to middle age in rats, and is associated with increased length, branching, and spine density of basal dendrites in CA1 hippocampal neurons. Neuroscience 202, 384–395, doi:10.1016/j.neuroscience.2011.11.053 (2012).

81 Retey, J. V. et al. A genetic variation in the adenosine A2A receptor gene (ADORA2A) contributes to individual sensitivity to caffeine effects on sleep. Clinical pharmacology and therapeutics 81, 692–698, doi:10.1038/sj.clpt.6100102 (2007).

82 Weibel, J. et al. Caffeine-dependent changes of sleep-wake regulation: evidence for adaptation after repeated intake. bioRxiv (2019).

