## Supplementary material for "Caffeine-induced Plasticity of Grey Matter Volume in Healthy Brains: A placebo-controlled multimodal within-subject study": Methods and supplements

Subject Recruitment and Participants

Applicants aged between 18 and 35 years, BMI ≥ 18 and ≤ 25, non-shift workers, and without a history of transmeridian travels <1 month prior to study, were screened according to the following exclusion criteria:

- self-reported caffeine intake <300 or >600 mg/day (calculations were based on the [1], adapted according to a classification of Snel and Lorist [2]) to ensure the safety of caffeine intake and to exclude extreme response,
- bad sleep quality, i.e. PSQI> 5 in the last four weeks assessed by the Pittsburg Sleep Quality Index (PSQI) to control for sleep disturbances
- extreme chronotype, as defined by Horne-Ostberg's Morningness-Eveningness Score (HOMES) ≤ 30 or ≥ 70 to prevent pronounced variance in circadian phase.
- self-reported regular substance use (including medication, nicotine, and drugs) and other major medical conditions.

A habituation night in the laboratory was conducted to exclude poor sleep efficiency (SE < 70%) and clinical sleep disturbances (apnea index > 10, periodic leg movements > 15/h). A toxicological screening right before each laboratory session served to control for drug intake including cannabis, amphetamine, methamphetamine, cocaine, benzocaine and morphine.

Participants, Study Protocol and Environmental Control

Overall, twenty healthy male volunteers completed both a 10-day caffeine and placebo condition. The average age was 26.4 ± 4.0 years, body mass index (BMI) 22.7 ± 1.38 kg/m2, and self-reported daily caffeine intake was 474.1 ± 107.5 mg/day.

Each of them underwent caffeine and placebo conditions, which each consisted of 9 days of ambulatory and 2 days of laboratory phases (Figure S1). In order to examine caffeine effects during daily intake as well as avoiding withdrawal effects [3] in the placebo condition, participants started caffeine (3 x 150 mg/day) or placebo (mannitol) capsules from the first day of ambulatory phase, i.e. 9 days before start of the laboratory phase. Timing of intake was set to 45 minutes, 4 hours, and 8 hours daily after waking up to model rather typical patterns of caffeine intake [4]. During the ambulatory phase, participants complied to a regular sleep-wake cycle (8 hours nighttime sleep, no naps allowed), ± 30 minutes in accordance with their habitual bedtime to control for accumulation of sleep debt, which was monitored and recorded by actimetry and self-reported sleep diary. Participants were asked to abstain from caffeine-containing diets including coffee, tea, energy drink, soda, and chocolate, etc., and their compliance was monitored by measuring the daily level of caffeine metabolites in fingertip sweat.

On the 9$^{th}$ day, participants started the laboratory phase, where they stayed in dim light (< 8 lux), constant half-supine position (~45°), with controlled dietary time and contents, as well as controlled toilet time. Access to the internet was forbidden. The participants slept at their habitual bedtime with polysomnographic recording. On the 10$^{th}$ day after waking up, the treatment (caffeine vs placebo) continued at identical times as during the ambulatory phase. The MRI scan took place at 4.5 hours after the third pill (by 12.5 hours of EEG-controlled wakefulness), in order to prevent the confounds of the acute effect of caffeine and reduce the influence of cerebral blood flow (CBF) on grey matter (GM) [5]. The time of the protocol was adapted to individual's habitual bedtime. Levels of caffeine and paraxanthine were measured in the fingertip sweat every 2 hours, and verbal working memory tasks (N-back) were performed every 4 hours through the laboratory phase.

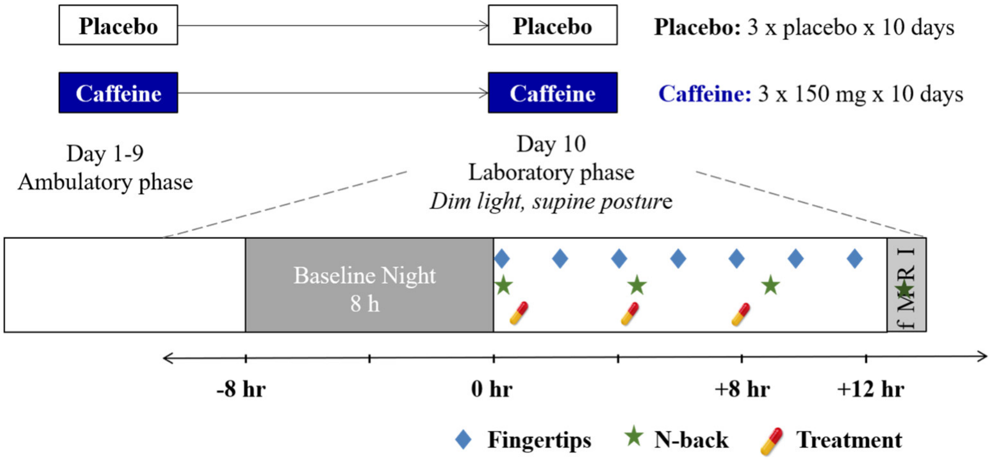


**Figure S1. Overview of the study design.**

Data acquisition

1. MRI

   T1-weighted structural data were obtained with MPRAGE sequence (1x1x1mm$^3$, TR=2000ms, TI= 1000ms, TE=3.37ms, FA=8°) on a 3T Siemens scanner (Prisma; Siemens AG, Erlangen, Germany). CBF was measured by 2D-EPI pulsed arterial spin labeling (ASL) sequence (4x4x4mm$^3$, TR=3000ms, TE=12ms, FA=90°, FoV=100) at the same MR scanner.

2. EEG

   Polysomnography was operated via V-Amp devices (Brain Products GmbH, Gilching, Germany) with a sampling rate of 500 Hz and notch filter at 50 Hz. The recording was conducted on the identical device between conditions of each subject. Electrophysiological activity was recorded above frontal (F3, F4), central (C3, C4), and occipital (O1, O2) regions, along with electro-oculography, electro-myography, and electro-cardiography to determine sleep stages. The current paper focuses on SWA (0.75- 4.5 Hz) derived from frontal regions in the first non-rapid eye movement (NREM) episode (defined by the period from the first epoch of NREM 1 stage until the last epoch of NREM 3 or 4 within the first sleep cycle), as this is known to be most sensitive for variations in sleep pressure [6-8].

3. Caffeine and metabolites from fingertip sweat

   The sample collection and the measurement were in collaboration with department of analytical chemistry, University of Vienna, together with Max F. Perutz Laboratories GmbH, Vienna, Austria. For the sample preparation, a 1 x 1 cm$^2$ filter papers wetted with water to collect the fingertip sweat were prepared by the Vienna laboratory. During sample collection, the experimenter first used alcohol wipes to clean up the participant's thumb and index finger of their non-dominant hands. Participants were told to hold tight the paper with two fingers for 1 minute, then the paper would be recollected back to a sealed tube. After collecting the samples, the caffeine and other metabolites were quantified by multiple-reaction monitoring protocol through the triple-quadrupole mass spectrometry.

4. N-back

We employed a visual N-back task with 4 hours interval through the day. For the current analysis we took the average of the 4 tasks from the morning to the last one in the scanner. To exclude habituation effects in the first session, participants performed one practice session in the evening before sleep. The task consisted of the visual presentation of a series of letters on a computer screen. Each task consisted of 9 trials of 3-back and 5 trials of 0-back, lasting in total approximately 15 minutes, whereby every trial contained 30 stimuli (letters), for each one must respond "yes" by keying down "1" or "no" by keying down "2". In 3-back trials, participants were asked to respond "yes" when the letter was identical to the letter three turns back while "no" for nonidentical stimuli. In 0-back trials, participants were asked to respond "yes" to each "K" letter while "no" for all others. In each trial, 10 stimuli correctly required a "Yes" response. Between the repetition of the tasks, the order of the trials and the consisted stimuli were designed in quasi-randomization in order to reduce a learning effect. The task was presented by the identical laptop device between conditions of each subject, and the brightness screen of the device was dimmed in order to control the interaction effect between light influence and caffeine [9].

Missing Data

**Table S1. Number of analyzed data sets along with missing.**

| Variable | N in Caffeine (ID of missing) | N in Placebo (ID of missing) |
|---|---|---|
| T1-weighted | 18 (#8, #13) | 19 (#3) |
| ASL | 17 (#8, #13, #15) | 17 (#3, #18, #20) |
| AUC-CAPX | 19 (#8) | 20 |
| SWA | 19 (#8) | 20 |
| N-Back | 17 (#8, #5, #16) | 19 (#16) |

* Reasons:

#8: Incompliant to treatment.

#3: Technical issue during fMRI data acquisition.

#13, #18, #20: Cut out part of sequences to catch up for delay in timing

#15: Technical issue during fMRI preprocessing.

#5, #16: Task misconducted (no rejection response)

Data Preprocessing and Analyses

1. Structural and CBF data

For the preprocessing of T1-weighted images, we used the CAT12 toolbox, running in SPM12 (University College London, London, UK). We adopted the built-in pipeline for longitudinal data (or repeated measures) where each volume of the subject was co-registered onto the mean of the three volumes. The average of each participant's total intracranial volume (TIV) and the volumes of each cerebral compartments (grey matter, white matter, and cerebrospinal fluid) were segmented by an affine registration on the SPM12 tissue probability map. The modulated normalization onto MNI space was carried out by first estimating one volumetric measure per subject which was then applied onto the other two repeated measures. A Gaussian kernel of FWHM= 8 x 8 x 8 $mm^3$ was adopted for the smoothing process.

ASL images were preprocessed with FMRIB Software Library (FSL) 5.0 developed by the Oxford Center for Functional MRI of the Brain (United Kingdom). Motion correction was estimated, followed by the generation of M0 calibration volume and tag-control pairs to calculate the relative and quantify the absolute CBF maps. The final absolute CBF maps that were co-registered onto the individual T1-weighted images and MNI space were ultimately employed in our subsequent multimodal analyses.

Volumetric differences in total GM between conditions were tested by a linear mixed model in R (R core team, Vienna, Austria). For voxel-based analyses of each single imaging modality, condition effects (caffeine vs placebo) and the covariation of SWA on regional GM and CBF were estimated by flexible factorial model in SPM12 and pair-T (equivalent to linear mixed model) on FSL, respectively.

Nonparametric threshold-free cluster enhancement (number of permutations = 5000, cluster-level threshold $p < .01$) was further performed by the SPM TFCE toolbox and FSL "*randomise*" function on GM and CBF, respectively. A mask of GM regions was applied to reduce the bias from the correction of multiple comparisons.

To test the covariation of sleep pressure with caffeine on GM changes, NREM sleep EEG SWA in the first sleep cycle was quantified as an index of sleep pressure. To examine the voxel-wise coefficient between multiple imaging modalities (i.e. CBF on GM), the VoxelStats toolbox [10] was applied to estimate the coefficient of CBF on GM differences with linear mixed model voxel by voxel (cluster-level threshold $p < .001$). Finally, we used VoxelStats to map out the covariation of both SWA and CBF with the effect of our treatment on GM. All the GM models were adjusted for individual total intracranial volumes (TIV).

2. EEG SWA

   The power density of SWA was quantified by Fast-Fourier-Transform spectrum analysis (T= 4s, window function = hamming, 0% overlapped). To determine NREM stage, two treatment-blinded experimenters (Y.-S. L., and J. W.) who have been trained and reached inter-rater reliability above 85% visually scored all nights in 30s epoch based on AASM [11].

3. Area under the curve of caffeine and paraxanthine levels

   We kept all values original including the ones below limit of detection. This does not affect the detection of a condition effect as the detection limits of caffeine (0.22 pg/FT) are very low. We used a combined value of caffeine (CA) and paraxanthine (PX), as paraxanthine also antagonizes to adenosine $A_1$, $A_{2A}$, and $A_{2B}$ receptors [12], while it follows a similar but slower metabolic pattern as compared to caffeine. To investigate the actual effect of acute caffeine exposure on other physical variables, the area under the curve (AUC) was calculated with the trapezoidal rule over the 7 samples from waking up til the scan. This measure is referred to as AUC-CAPX.

4. N-Back

The mean rates of four types of responses (hit, missed, false alarm, and correct rejection) were defined as the proportion of the responses out of total targets within one task. Correct rate was defined as (hits + correct rejection)/all responses + no response, while incorrect rate was (missed + false alarm)/all responses + no response. Accuracy was defined as the ratio of correct rate to incorrect rate. The mean reaction time of overall as well as of each type of responses was calculated over 9 bouts of each task. From waking up until the scan, participants accomplished 4 times of tasks every 4 hour, the mean values of accuracy and reaction time over the 4 times were used to modeling.

Model Details

Table S2 summarizes the models adopted on R to estimate the condition effect, together with the covariates (CBF and SWA), on the three main dependent variables (DV) – *total* GM, mTL GM, and N-back parameters. The data points of the regional GM volumes were the eigenvariate extracted from the thresholded clusters expressing a significant caffeine effect. For CBF and SWA, Baron and Kenny's step-wise models [13] were utilized to examine the mediation on caffeine-associated GM changes. The choice of the distribution of the models was based on the distribution, the type of responses of the DVs, as well as the tests on their goodness of fit.

All models including total or regional GM were adjusted for the individual total intracranial volume (TIV) estimated in each scan. All estimations have been adjusted for the order of the study to control for learning effects.

**Table S2. A list of adopted parameters in the linear and generalized linear mixed models.**

| DV | Main effect (fixed) | Covariates (fixed) | Random effect | Method | Family(link) |
|---|---|---|---|---|---|
| TGM | Condition[a1] | TIV | Subject | GLMM | Gamma (log) |
| | AUC-CAPX [a2] | TIV | Subject<br>$\beta_0$ + AUC-CAPX * Condition | | |
| | CBF [a3] | TIV | Subject<br>$\beta_0$ + CBF * Condition | | |
| | Condition * SWA [a4] | TIV | Subject<br>$\beta_0$ + Condition | | |
| | Condition [a1] | TIV<br>SWA or CBF | Subject | | |
| mTL GM | Condition [b1] | TIV | Subject | LMM | Gaussian |
| | log (AUC-CAPX) [b2] | TIV | Subject<br>$\beta_0$ + AUC-CAPX * Condition | | |
| | CBF [b3] | TIV | Subject<br>$\beta_0$ + CBF * Condition | | |
| | SWA [b4] | TIV | Subject<br>$\beta_0$ + Condition | | |
| | Condition [a1] | TIV<br>SWA or CBF | Subject | | |
| Accuracy in 3-N | Condition [a1] | 0-N accuracy<br>Study order | Subject<br>Experimental order | GLMM | Gamma (log) |

| | | | | | |
|---|---|---|---|---|---|
| | AUC-CAPX [a1] | 0-N accuracy<br>Study order | Subject<br>Experimental order<br>AUC-CAPX ~ $\beta_0$ + Condition | | |
| RT<br>in 3-N | Condition [a1] | 0-N RT<br>Study order | Subject<br>Experimental order | GLMM | Gamma (log) |
| | AUC-CAPX [a1] | 0-N RT<br>Study order | Subject<br>Experimental order<br>AUC-CAPX ~ $\beta_0$ + Condition | | |

Supplemented Results

1. CAPX levels across time

The average summation of caffeine and paraxanthine overall was 10.20 ± 13.83 ng/filter in the placebo condition and 120.79 ± 77.85 ng/filter in caffeine condition.

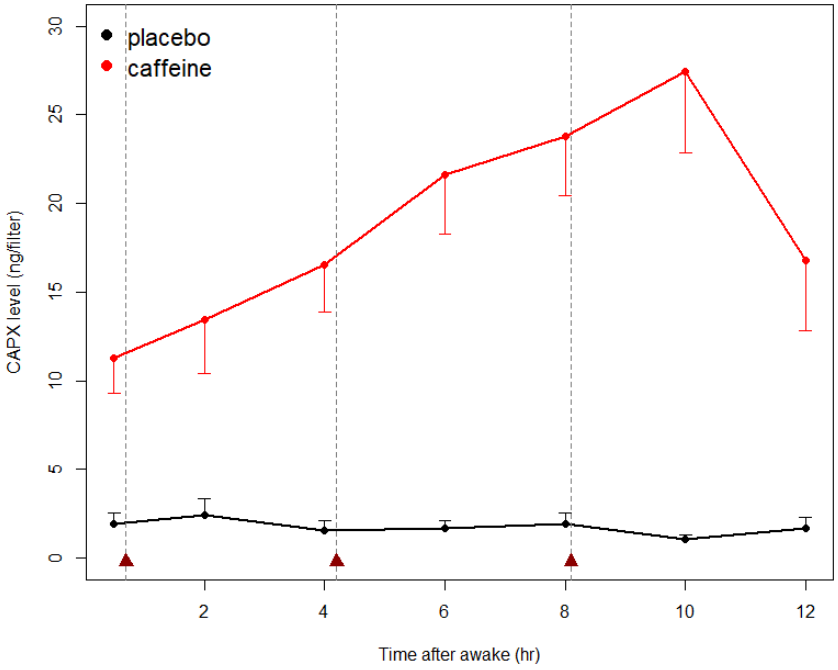


**Figure S2. The dynamic of group average AUC-CAPX across the laboratory day (7 timepoints from the awakening to 30 minutes before the MRI scan).** The 3 timepoints of treatment administration are presented in triangle

2. NREM SWA power density

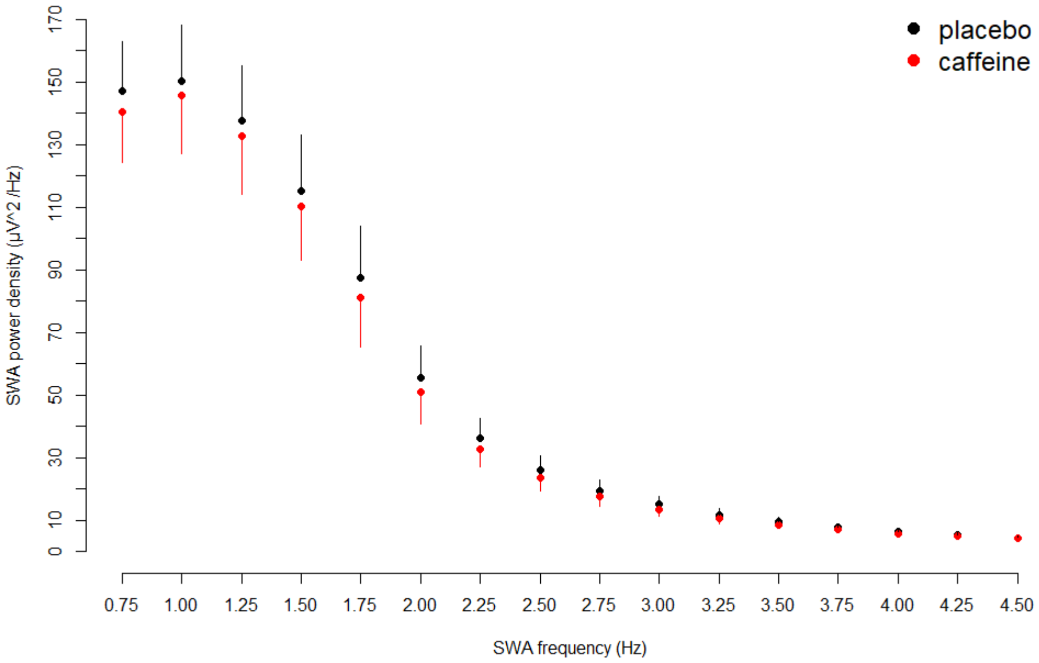


**Figure S3. Group average power density of slow-wave activity in 0.25 bin during the first NREM cycle.** Average log-transformed power density of overall SWA frequency band is 3.05 ± 0.67 μV^2 /Hz in placebo condition and 2.96 ± 0.70 μV^2 /Hz in caffeine condition.

3. N-Back Tasks

The inspection of each type of response indicated that volunteers tended to have more incorrect responses (missed and false alarm) and less correct responses (hits and correct rejection) in the caffeine condition, however only the type of false alarm reached statistical significance ($t_{con}$ = 45.20, $p_{con}$ < .001; $R^2m$ = .20; $R^2c$ = .69). On the other hand, the RT in different types of responses exhibited distinguished changes after long-term caffeine intake.

**Table S3. Condition effects on each parameter of working memory performance and its association with AUC-CAPX.**

| | **Condition (Caff vs Plac)** | **(Higher) AUC-CAPX** |
|---|---|---|
| Accuracy 0-N | Worse<br>95% CI= [ -0.041, -0.004]<br>$R^2m$ = 0.03; $R^2c$ = 0.32 | N.S. |
| Accuracy 3-N | Worse<br>95% CI= [-0.073, -0.001]<br>$R^2m$ = 0.03; $R^2c$ = 0.57 | N.S. |
| Accuracy 3-N controlling for 0-N | Worse<br>95% CI= [ -0.036, > -0.001]<br>$R^2m$ = 0.41; $R^2c$ = 0.85 | Better<br>95% CI= [0.002, 0.008]<br>$R^2m$ = 0.87; $R^2c$ = 0.99 |
| Hits (mean rate) * | N.S. | Better<br>95% CI= [0.006, 0.026]<br>$R^2m$ = 0.87; $R^2c$ = 0.99 |
| Correct rejection (mean rate) * | N.S. | Better<br>95% CI= [0.002, 0.006]<br>$R^2m$ = 0.92; $R^2c$ = 0.99 |
| Missed (mean rate) * | N.S. | N.S. |
| False alarm (mean rate) * | Worse<br>95% CI= [0.103, 0.113]<br>$R^2m$ = 0.20; $R^2c$ = 0.69 | N.S. |
| RT (Hits) * | N.S. | Faster<br>95% CI= [-0.029, -0.008]<br>$R^2m$ = 0.90; $R^2c$ = 0.99 |
| RT (Correct Rejection) * | Slower<br>95% CI= [> -0.001, 0.136]<br>$R^2m$ = 0.13; $R^2c$ = 0.66 | Faster<br>95% CI= [-0.032, -0.007]<br>$R^2m$ = 0.90; $R^2c$ = 0.99 |

| | | |
|---|---|---|
| RT (Missed) * | Slower<br>95% CI= [0.037, 0.219]<br>$R^2m = 0.05$; $R^2c = 0.68$ | Faster<br>95% CI= [-0.037, -0.004]<br>$R^2m = 0.87$; $R^2c = 0.99$ |
| RT (False Alarm) * | N.S. | Faster<br>95% CI= [-0.040, -0.015]<br>$R^2m = 0.91$; $R^2c = 0.99$ |

* Responses in 3-back controlling for 0-back.

1 Bühler, E., Lachenmeier, D. W., Schlegel, K. & Winkler, G. Development of a tool to assess the caffeine intake among teenagers and young adults. *Science and Research* **61**, 58-63, doi: 10.4455/eu.2014.011 (2013).
2 Snel, J. & Lorist, M. M. Effects of caffeine on sleep and cognition. *Progress in brain research* **190**, 105-117, doi:10.1016/b978-0-444-53817-8.00006-2 (2011).
3 Juliano, L. M. & Griffiths, R. R. A critical review of caffeine withdrawal: empirical validation of symptoms and signs, incidence, severity, and associated features. *Psychopharmacology* **176**, 1-29, doi:10.1007/s00213-004-2000-x (2004).
4 Martyn, D., Lau, A., Richardson, P. & Roberts, A. Temporal patterns of caffeine intake in the United States. *Food Chem. Toxicol.* **111**, 71-83, doi:10.1016/j.fct.2017.10.059 (2018).
5 Couturier, E. G., Laman, D. M., van Duijn, M. A. & van Duijn, H. Influence of caffeine and caffeine withdrawal on headache and cerebral blood flow velocities. *Cephalalgia : an international journal of headache* **17**, 188-190, doi:10.1046/j.1468-2982.1997.1703188.x (1997).
6 Cajochen, C., Khalsa, S. B., Wyatt, J. K., Czeisler, C. A. & Dijk, D. J. EEG and ocular correlates of circadian melatonin phase and human performance decrements during sleep loss. *The American journal of physiology* **277**, R640-649 (1999).
7 Finelli, L. A., Baumann, H., Borbely, A. A. & Achermann, P. Dual electroencephalogram markers of human sleep homeostasis: correlation between theta activity in waking and slow-wave activity in sleep. *Neuroscience* **101**, 523-529 (2000).
8 Werth, E., Achermann, P. & Borbely, A. A. Fronto-occipital EEG power gradients in human sleep. *Journal of sleep research* **6**, 102-112 (1997).
9 Wright, K. P., Jr., Badia, P., Myers, B. L., Plenzler, S. C. & Hakel, M. Caffeine and light effects on nighttime melatonin and temperature levels in sleep-deprived humans. *Brain research* **747**, 78-84 (1997).
10 Mathotaarachchi, S. *et al.* VoxelStats: A MATLAB Package for Multi-Modal Voxel-Wise Brain Image Analysis. *Frontiers in neuroinformatics* **10**, 20, doi:10.3389/fninf.2016.00020 (2016).
11 Berry, R. B. *et al.* The AASM Manual for the Scoring of Sleep and Associated Events: Rules, Terminology and Technical Specifications, Version 2.0. *Darien, Illinois: American Academy of Sleep Medicine* (2012).
12 Cornelis, M. C. *et al.* Genome-wide association study of caffeine metabolites provides new insights to caffeine metabolism and dietary caffeine-consumption behavior. *Hum. Mol. Genet.* **25**, 5472-5482, doi:10.1093/hmg/ddw334 (2016).
13 Baron, R. M. & Kenny, D. A. The moderator-mediator variable distinction in social psychological research: conceptual, strategic, and statistical considerations. *Journal of personality and social psychology* **51**, 1173-1182 (1986).
